# Increased Histone-DNA Complexes and Endothelial-Dependent Thrombin Generation in Severe COVID-19

**DOI:** 10.1101/2021.07.03.450992

**Authors:** Beth A. Bouchard, Christos Colovos, Michael Lawson, Zachary Osborn, Adrian Sackheim, Kara J. Mould, William J. Janssen, Mitchell J. Cohen, Devdoot Majumdar, Kalev Freeman

## Abstract

**Objective:** Coagulopathy in severe COVID-19 is common but poorly understood. The purpose of this study was to determine how SARS-CoV-2 infection impacts histone levels, fibrin structure, and endogenous thrombin potential in the presence and absence of endothelial cells.

**Approach:** We studied individuals with SARS-CoV-2 infection and acute respiratory distress syndrome at the time of initiation of mechanical ventilation compared to healthy controls. Blood samples were assayed for levels of histone-DNA complexes. Confocal microscopy was used to evaluate fibrin structure in clots formed from recalcified plasma samples using fluorescently-labeled fibrinogen. Endogenous thrombin potential was measured by calibrated automated thrombin assays in the presence of tissue factor and phospholipid (PCPS) or cultured human endothelial cells.

**Results:** Circulating nucleosomes were elevated in the plasma of COVID-19 patients relative to healthy controls (n=6, each group). COVID-19 patient plasma thrombin generation was also altered. Despite having an increased endogenous thrombin potential, patient plasma samples exhibited prolonged lag times and times to peak thrombin in the presence of added tissue factor and PCPS. Strikingly different results were observed when endothelial cells were used in place of tissue factor and PCPS. Control plasma samples did not generate measurable thrombin (lag time >60 min); in contrast, plasma samples from COVID-19+ patients generated thrombin (mean lag time ∼20 min). Consistent with the observed alterations in thrombin generation, clots from COVID-19 subjects exhibited a denser fibrin network, thinner fibers and lower fibrin resolvability.

**Conclusions:** Elevated histones, aberrant fibrin formation, and increased endothelial-dependent thrombin generation in COVID-19 may contribute to coagulopathy.

**HIGHLIGHTS:**

- Histone-DNA complexes are significantly elevated in the plasma of patients with severe SARS-CoV-2 infection.
- Measures of thrombin generation by calibrated automated thrombography and fibrin clots formed *in situ* are altered in severe COVID-19.
- Plasma from COVID-19 patients promotes thrombin generation on cultured endothelial cells in the absence of added tissue factor or phospholipids.
- The additive effects of histones on thrombin generation and endothelial cell function may play a major role in the thrombotic complications observed in severe SARS-CoV-2 infection.

## INTRODUCTION

The emergence of the severe acute respiratory syndrome novel corona virus 2 (SARS-CoV-2) resulted in a global pandemic that afflicted over 182 million people globally and over 33 million in the United States alone as of July 2021. The associated disease, coronavirus disease 2019 (COVID-19), accounts for nearly 4 million deaths worldwide and over 590,000 in the United States [1]. A consistent finding amongst patients with SARS-CoV-2 is derangements in coagulation markers and increased incidence of thrombotic complications [2-6]. Derangement in coagulation parameters was associated with a coagulation phenotype similar to disseminated intravascular coagulopathy and associated with death [5, 7]. Findings of micro and macro vascular thromboses resulting in organ failure are observed in autopsy studies. Emerging evidence of the benefits provided by therapeutic anticoagulation further supports the idea that coagulopathy contributes to SARS-CoV-2 disease progression.

Cellular release of histones, major mediators of death in sepsis and neutrophil extracellular trap (NET) formation, has been implicated in COVID-19 coagulopathy [8-12]. Histones increase plasma thrombin generation directly through interactions with coagulation proteins [13, 14], platelets, neutrophils, and endothelial cells [15, 16]. The purpose of this study was to test the hypothesis that elevations in circulating histones during SARS-CoV-2 infection contribute to coagulopathy by determining how SARS-CoV-2 infection impacts histone levels, fibrin structure, and thrombin generation in the presence and absence of endothelial cells.

## METHODS

### Study participants

We recruited hospitalized ICU patients (n=9) with acute SARS-CoV-2 infections confirmed by RT-PCR at St. Joseph’s Hospital in Denver, CO. All patients required mechanical ventilation due to the severity of lung infection. All participants provided written consent using a protocol approved by National Jewish Health IRB. Healthy donors (n=6) were recruited and provided written consented using a protocol approved by the University of Vermont Committee on Human Research.

### Sample acquisition

Plasma from patients and healthy donors was prepared from citrated (3.2%) whole blood obtained at the onset of mechanical ventilation from a central line or the antecubital vein using standard methods. All plasma samples were stored at -80°C in small aliquots.

### Measurement of histone levels

Cell-free histone levels were assessed in plasma from healthy controls and COVID-19 patients using a photometric enzyme immunoassay (Cell Death Detection ELISAPLUS kit, Roche Applied Science) that measures histone-associated DNA fragments according to the manufacturer’s instructions.

### In situ fibrin polymerization

Fibrin clot formation was assessed as previously described [17]. Plasma was diluted 1:1 with HBS and incubated on glass chamber slides with AlexaFluor488-labeled fibrinogen (220 nM) (ThermoFisher Scientific, Waltham, MA), tissue factor (8.7 pM) and CaCl_2_ with HistoChoice (MilliporeSigma, Burlington, MA), and an anti-photobleaching agent (Agilent Technologies, Santa Clara, California). Clots were visualized by confocal microscopy using a Nikon A1R Confocal Microscope (Nikon Instruments Inc, Melville NY) with a 60X/1.5 oil immersion objective. For each clot, a three-dimensional “Z-stack” image series consisting of 40 images at 0.25 µm steps through each sample was obtained. Plasma clots were formed and imaged in duplicate.

### Endothelial cell culture

Human endothelial cells (EA.hy926; ATCC® CRL-2922™) were cultured in Dulbecco’s Modified Eagle Medium (DMEM) supplemented with 10% fetal bovine serum and 5 µg/mL gentamycin (complete medium) at 37°C, 5% CO_2_. Prior to use, the complete medium was removed and the cells incubated with DMEM for 1 hr at 37°C, 5% CO_2_. The cells were released from the tissue culture wells with trypsin and subjected to centrifugation (170 x g, 7 min). Cell pellets were washed one time by resuspension in 20 mM HEPES, 0.15 M NaCl (pH 7.4) (HBS) followed by centrifugation. The final cell pellets were resuspended in HBS and adjusted to a final concentration of 1×10^7^/mL.

### Thrombin generation

Thrombin generation was assessed by a modified calibrated automated thrombography (CAT) [18]. Plasma was thawed at 37°C in the presence of corn trypsin inhibitor (0.1 mg/mL final concentration). Plasma was incubated with the thrombin substrate Z-Gly-Gly-Arg 7-amido-4-methylcoumarin hydrochloride (0.42 mM) (Bachem AG, Switzerland) and CaCl_2_ (15 mM) (3 min, 37°C), and the reactions initiated by the addition of relipidated tissue factor_1-242_ (6.5 pM) (a gift from Dr. R. Lunblad, Baxter Healthcare Corp.) and synthetic vesicles consisting of 80% phosphatidylcholine and 20% phosphatidylserine (PCPS) (20 µM), or EA.hy926 cells (2×10^5^). Fluorescence was measured (ex = 370 nm/em = 460 nm) for 1 hour with a Cytation 3 imaging reader (BioTek, Winooski, VT). Changes in fluorescence were converted to thrombin concentrations using a calibration curve created from sequential dilutions of human thrombin. If no change in fluorescence was noted after 60 min, the lag time for the sample was defined as >60 min.

### Statistical analysis

Data are expressed as mean ± standard error of the mean. Unpaired, 2-tailed, t-tests were performed to compare the lag times and mean histone levels between controls and patients. For comparison of clot resolvability, the Kolmogorov-Smirnov test was used to compare cumulative distributions with 95% confidence. A p-value less than 0.05 was considered significant.

## RESULTS

We enrolled patients diagnosed with COVID-19 (n=9 patients) and healthy controls (n=6). The patient characteristics are described in Table 1. Three patients who had lag times >60 min in thrombin generation assays in the presence of added tissue factor and PCPS due to therapeutic anticoagulation were excluded from analysis.

**Table 1.** COVID-19 patient characteristics.

| Patient | Survival<br>(day 28) | Age<br>(years) | Gender | Race/<br>Ethnicity | BMI* | Diabetes | HTN† |
| --- | --- | --- | --- | --- | --- | --- | --- |
| 1 | Y | 53 | M | White | 30.8 | Y | Y |
| 2 | Y | 54 | M | Hispanic or<br>Latino | 30.4 | N | N |
| 3 | Y | 57 | F | White | 28.6 | Y | Y |
| 4 | Y | 68 | F | Hispanic or<br>Latino | 38.9 | N | N |
| 5 | Y | 69 | F | Hispanic or<br>Latino | 33.6 | Y | Y |
| 6 | Y | 70 | M | Hispanic or<br>Latino | 32.0 | N | Y |
| 7 | N | 62 | M | Black | 24.8 | N | Y |
| 8 | N | 63 | F | Black | 27.5 | N | Y |
| 9 | N | 64 | M | White | 41.1 | Y | N |

\* Body mass index, kg/m<sup>2</sup>
† Hypertension

Table 1 (cont.). COVID-19 patient characteristics.
| Patient | Smoker | Steroids | Convalescent plasma | RDV<sup>‡</sup> | CMV<sup>§</sup> (days) | CMV (total days) | Days since admit |
| --- | --- | --- | --- | --- | --- | --- | --- |
| 1 | Never | N | N | N | 2 | 17 | 4 |
| 2 | Never | Y | N | N | 6 | 19 | 6 |
| 3 | Never | Y | Y | Y | 7 | 22 | 13 |
| 4 | Former | Y | Y | Y | 4 | 36 | 7 |
| 5 | Never | N | N | N | 3 | 8 | 5 |
| 6 | Never | Y | Y | Y | 4 | 7 | 4 |
| 7 | Current | N | N | N | 8 | 15 | 8 |
| 8 | Former | N | N | Y | 4 | 24 | 4 |
| 9 | Current | Y | Y | Y | 5 | 7 | 7 |

<sup>‡</sup> Remdesivar
<sup>§</sup> Continuous mandatory ventilation

Table 1 (cont.). COVID-19 patient characteristics.
| Patient | Days since<br>+ COVID<br>test | Arterial pH | P/F ratio | APACHE<br>score | SAPS<sup>#</sup> | LIPS<sup>**</sup> |
| --- | --- | --- | --- | --- | --- | --- |
| 1 | 7 | 7.52 | 443 | 16 | 35 | 5.5 |
| 2 | 6 | 7.48 | 134 | 10 | 37 | 6.5 |
| 3 | 17 | 7.41 | 166 | 16 | 41 | 3.5 |
| 4 | 6 | 7.35 | 94 | 14 | 41 | 6.5 |
| 5 | 11 | 7.44 | 220 | 17 | 36 | 7 |
| 6 | 4 | 7.44 | 150 | 15 | 43 | 4.5 |
| 7 | 8 | 7.31 | 125 | 17 | 51 | 7 |
| 8 | 4 | 7.45 | 262 | 11 | 37 | 5.5 |
| 9 | 7 | 7.26 | 159 | 21 | 51 | 7.5 |

<sup>||</sup> Acute physiologic assessment and chronic health evaluation
<sup>#</sup> Simplified acute physiology score
<sup>\*\*</sup> Lung injury prediction score

Using an ELISA that detects cell-free histone-associated DNA fragments, it was observed that histones were significantly elevated in COVID-19 patient plasma relative to plasma from healthy controls (Figure 1A).

**Figure 1.**
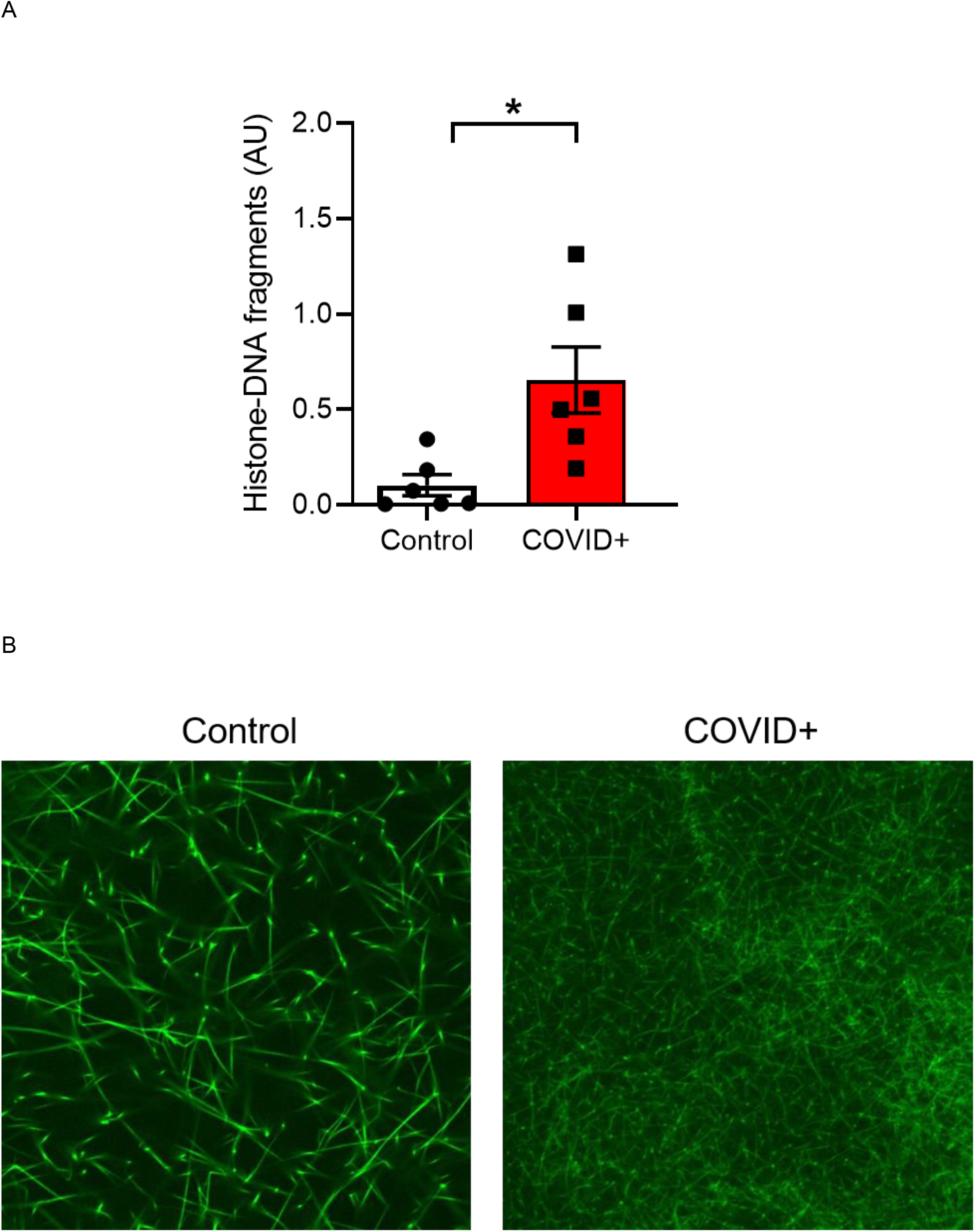
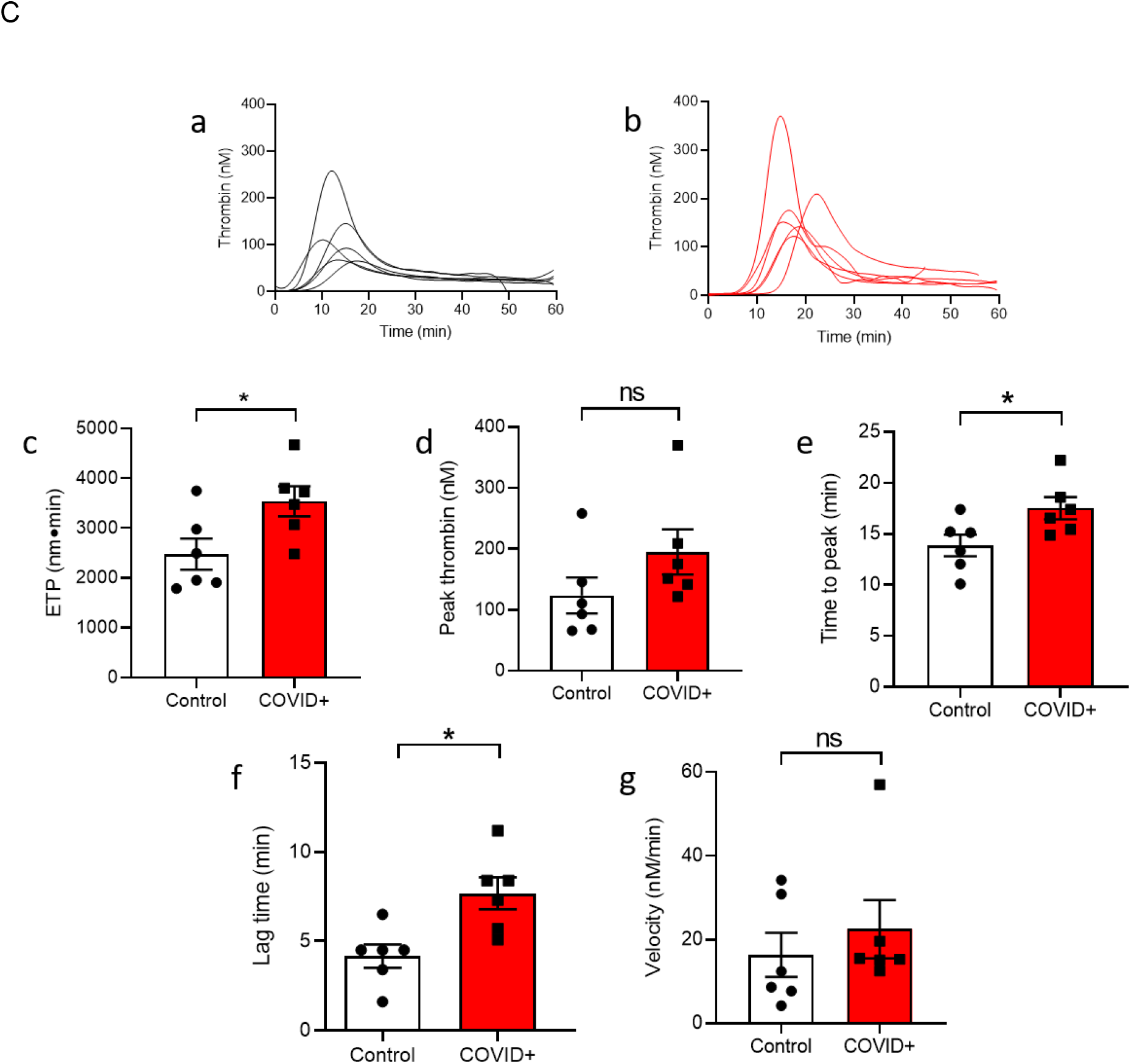
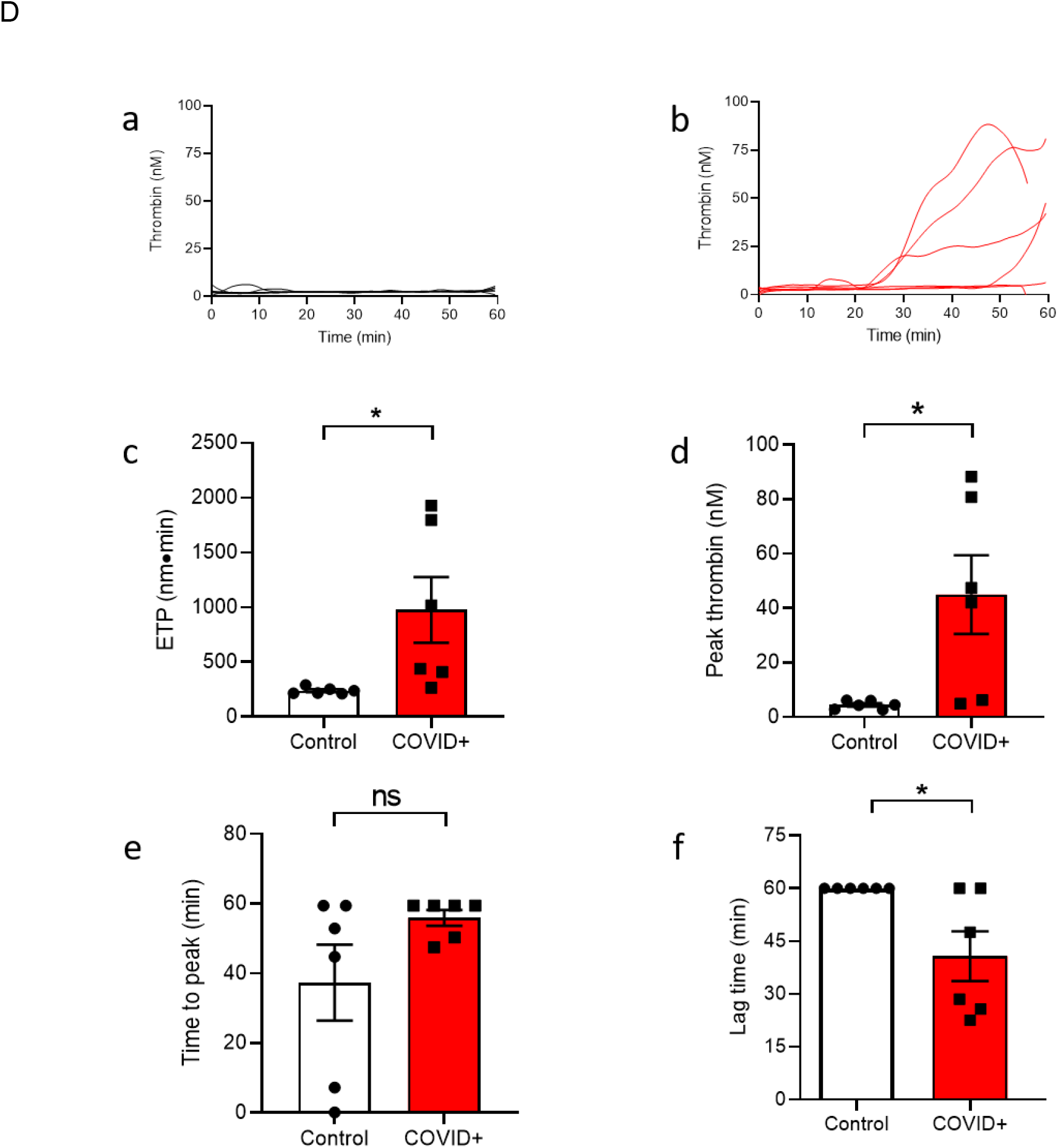
Histone levels, fibrin clot structure and endogenous thrombin potential are altered in plasma from COVID-19 patients. (A) Histone-DNA fragments were measured in healthy controls (n=6) and COVID-19 patients (n=6) using a commercially available assay (*P=0.0126). (B) Representative confocal microscopy images of fluorescently labeled fibrinogen polymerized in the presence of control (*left*) and COVID+ patient (*right*) plasma samples. Clots were imaged on a Nikon A1R Confocal microscope (60X/1.5 oil immersion). Each image is a representative cross-section from a three-dimensional Z-stack (0.25 µm steps) of 40 images captured from each clot. Calibrated automated thrombography was used to measure thrombin generation using plasma from healthy controls (n=6) or COVID-19 patients (n=6) in the presence of (C) tissue factor and PCPS or (D) EA.hy926 cells. The mean of duplicate thrombin generation curves for each healthy control (a) and COVID-19 patient (b) is shown. Endogenous thrombin potential (c), peak thrombin (d), time to peak thrombin (e), lag time (f) and rate of thrombin generation (velocity) (g) of the controls and patients under these conditions were compared. *p<0.05 was considered significant. For the plasma samples that did not clot in the presence of cells over the duration of the experiment (60 min), the lag times were set to 60 min. ETP = endogenous thrombin potential; ns = not significant.

Fibrin clot structure was also assessed using AlexaFluor488-labeled fibrinogen. Comparison of the clot structures by confocal microscopy demonstrated that fibrin clots formed using plasma from COVID-19 patients were markedly different from those formed using healthy control plasma with a denser fibrin network, thinner fibers and lower fibrin resolvability (Figure 1B) suggesting that thrombin generation in the COVID-19 plasma was altered (PMID: 17208341).

Thrombin generation in healthy control plasma and plasma from COVID-19 patients were compared by CAT in the presence of added tissue factor and PCPS (Figure 1C, panels a & b) or cultured EA.hy926 cells (Figure 1D, panels a & b). In the presence of tissue factor and PCPS, the endogenous thrombin potential (panel c) of plasma from COVID-19 patients (n=6) was significantly higher than plasma from healthy controls (n=6) although peak levels of thrombin (panel e) were not different. Despite the apparent hypercoagulability of the COVID-19 patient plasma, significantly longer lag times (panel e) and times to peak thrombin (panel f) were observed in plasma from patients as compared to healthy controls, and the rates of thrombin generation (panel g) were not different. Strikingly different results were observed when EA.hy926 cells were used in place of added tissue factor and PCPS (Figure 1D). Under these conditions, none of the healthy control plasma samples generated thrombin (lag time >60 min) (panels a & e). Unexpectedly, and in marked contrast to the results obtained with control plasma, four of the COVID-19 patient plasma samples generated thrombin (mean lag time ∼20 min) (panels b & e) albeit at levels lower than that observed in the presence of added tissue factor and PCPS. Consistent with this observation, the mean endogenous thrombin potential (panel c) and peak levels of thrombin (panel d) were significantly higher in patient vs healthy control plasma, while the times to peak thrombin (panel f) were not different. Rates of thrombin generation were unable to be calculated in the absence of added tissue factor and PCPS. These combined observations suggest that thrombin generation by plasma from patients with severe COVID-19 infections is dysregulated, and that high levels of histones or another molecule(s) effect(s) blood coagulation in the presence of endothelial cells.

## DISCUSSION

Histones act as damage-associated molecular pattern molecules following their release from cells by NETs, cell apoptosis, or cell necrosis [19]. Circulating levels of nucleosomes and histones are significantly elevated and correlate with the severity or poor outcome of several pathophysiological processes such as acute bacterial infection, sepsis, autoimmune diseases, cerebral stroke, trauma, cancer [19], and acute pulmonary embolism [20]. Here, we show increased histone-associated DNA fragments in SARS-CoV-2 infection resulting in severe COVID-19. This is consistent with prior reports of elevated NETs in COVID-19, as evidenced by increased cell-free DNA, myeloperoxidase-DNA complexes, and citrullinated-Histone H3 [21-23]. Similar observations were recently made in two independent cohorts of COVID-19 positive patients with a quantitative nucleosome immunoassay that measured cell-free H3.1 nucleosomes [24]. These investigators demonstrated that nucleosomes were highly elevated in plasma of COVID-19 patients with a severe course of the disease relative to healthy controls and that both the histone 3.1 variant and citrullinated nucleosomes increase with disease severity.

Extracellular histones enhance plasma thrombin generation by reducing thrombomodulin-dependent protein C activation [14]. Additionally, isolated histone proteins bind to prothrombin via fragment 1 and fragment 2 (non-catalytic portions), and reduce the need for factor Xa in clotting [13]. Consistent with our observations are very recent studies by Bouck and colleagues [25] who also demonstrated that compared with healthy donors, patients with SARS-CoV-2 infection had increased thrombin generation potential but an unexpectedly prolonged lag time. Additional experiments demonstrated increased endogenous plasmin potential and delayed plasmin formation. These perturbations led to increased fibrin formation, and combined with our study, support the idea that histones and/or NETs contribute to the observed dysregulation of hemostasis in severe COVID-19.

The fibrin clots formed *in situ* by plasma from patients with severe COVID-19 are denser with thinner fibers as compared to clots formed from the plasma of healthy controls. Patients with thromboembolic diseases also form structurally abnormal clots that are resistant to fibrinolysis [26]. Similar observations have been made in patients with severe trauma [17], which is also characterized by a disseminated intravascular coagulation-like phenotype [27]. As patients with acute thromboembolic disease [20] and trauma [13] also have elevated circulating histone levels, these structurally- and functionally-altered clots may be the result of covalent (via factor XIIIa crosslinking) and noncovalent interactions of histones with fibrin [28] in addition to dysregulated thrombin formation.

Histones can also promote thrombotic events by inducing endothelial cells to release pro-inflammatory cytokines [15], increase cell surface adhesion molecules [16], and express tissue factor [15]. When cultured endothelial cells were used in place of added tissue factor and PCPS, we observed considerable thrombin generation by COVID-19 patient plasma suggesting that histones or (an)other molecule(s) present in COVID-19 plasma induce(s) an inflammatory and procoagulant endothelial cell phenotype. In support of this notion is the observation that post-mortem analyses of lung samples from patients with COVID-19 demonstrate higher tissue expression of interleukin-6, tumor necrosis factor-α, intracellular adhesion molecule-1 and caspase-1, a marker of pyroptosis, an inflammatory form of programmed cell death, in lung endothelial cells as compared to H1N1 and control groups [29]. Additional studies with a larger patient cohort and access to larger plasma volumes are warranted to test these hypotheses. In conclusion, the results of this small study demonstrate that circulating histones are elevated in severe COVID-19 and suggest that they can influence thrombin formation through one or more mechanisms. The combined effects of histones may have profound effects on thrombin generation, especially in the presence of endothelial cells, and play a major role in COVID-19 coagulopathy. Neutralization of histone effects may mitigate these coagulopathic responses and reduce morbidity and mortality.

## ACKNOWLEDGMENTS

The authors would like to acknowledge the generous donation of recombinant tissue factor_1-242_ by Dr. R. Lunblad, Baxter Healthcare Corp.

